# A Conserved Flexible N-Terminal Domain Tunes the Calcium Sensitivity of Sorcin by Stabilizing Its Active Conformation

**DOI:** 10.64898/2026.07.29.741579

**Authors:** Qiushi Ye, Angela Wu, Kathleen Joyce Carillo, Ibrahim D. Boyenle, Holly Hemesath, Yanan He, Nicolas Delaeter, Jaekyun Jeon, John Orban, Lei Zhang, Yanxin Liu

## Abstract

Sorcin is a penta-EF-hand Ca^2+^-binding protein that acts as a Ca^2+^ sensor and regulator of Ca^2+^ homeostasis. Although the structure and Ca^2+^-dependent activation of Sorcin is well characterized, the function of its flexible N-terminal domain (NTD) remains unclear. We combined sequence analysis, Ca^2+^-induced aggregation assays, multidimensional NMR spectroscopy, and long-timescale molecular dynamics (MD) simulations to define the NTD’s role in Sorcin activation. Sequence comparisons showed that the NTD is conserved across vertebrates despite its intrinsic disorder, indicating functional importance. NTD deletion markedly reduced Ca^2+^ responsiveness. Relative to full-length Sorcin, the construct with NTD truncation required more than twofold higher Ca^2+^ concentrations and over tenfold higher protein concentrations to initiate aggregation, while aggregation kinetics slowed by nearly three orders of magnitude. Temperature-dependent measurements yielded an apparent activation energy of 36.7 kJ·mol^-1^, consistent with aggregation driven by Ca^2+^-induced conformational activation rather than denaturation. NMR chemical shift perturbations localized the effects of NTD removal to the EF-hand Ca^2+^-binding loops and adjacent D-helix. Apo-state MD simulations revealed transient intra- and intermolecular NTD-SCBD contacts that explain some perturbations and support direct and allosteric regulation. Ca^2+^-bound simulations further showed that SCBD departs more readily from the crystallographic active conformation than full-length Sorcin, indicating that the NTD stabilizes the active state through dynamic contacts. Together, these findings establish the NTD as a critical regulator that enhances Ca^2+^ responsiveness by shifting Sorcin’s conformational equilibrium toward the active state and suggest that flexible N-terminal extensions regulate signaling within the penta-EF-hand protein family.

## Introduction

Calcium ions (Ca^2+^) function as universal second messengers that regulate a wide range of cellular processes, including muscle contraction, secretion, metabolism, gene expression, and apoptosis^1–4^. The specificity of Ca^2+^ signaling is achieved through a diverse array of Ca^2+^-binding proteins that sense changes in intracellular Ca^2+^ concentration and convert these signals into downstream biological responses^5,6^. Among these proteins, EF-hand Ca^2+^-binding proteins represent one of the largest and most extensively studied families of Ca^2+^ sensors^7–9^. While many EF-hand proteins respond to submicromolar or low-micromolar Ca^2+^ concentrations, others are adapted to detect the higher Ca^2+^ levels that occur within specialized cellular microdomains^6,10,11^.

Sorcin (soluble resistance-related calcium-binding protein, SRI) is a highly conserved member of the penta-EF-hand (PEF) protein family^12–14^. Originally identified in multidrug-resistant cancer cells^15,16^, Sorcin is now recognized as an important regulator of intracellular Ca^2+^ homeostasis through its interactions with Ca^2+^ channels, pumps, and transporters^13,17–19^. Sorcin is particularly abundant in excitable tissues such as the heart and plays critical roles in excitation-contraction coupling, endoplasmic reticulum/sarcoplasmic reticulum Ca^2+^ handling, and cellular stress responses^20–22^. Dysregulation of Sorcin has been implicated in cancer^23,24^, cardiac dysfunction^25^, and neurodegenerative diseases^26^, underscoring its physiological and biomedical importance^13,18^.

Structurally, Sorcin forms a homodimer consisting of a conserved C-terminal calcium-binding domain (SCBD) containing five EF-hand motifs and a glycine-rich N-terminal domain (NTD) (Figure 1A, left)^12,14,27^. Crystal structures have captured both apo and Ca^2+^-bound conformations of Sorcin and revealed substantial rearrangements of the calcium-binding domain upon Ca^2+^ binding^12,14,28^. These conformational changes expose hydrophobic surfaces that facilitate interactions with target proteins and promote Sorcin self-association^11,12,14,28^. Notably, in the Ca^2+^- bound structure a segment of Sorcin’s own conserved, glycine-rich N-terminal motif was observed bound within one of these exposed hydrophobic pockets (Figure 1A, right)^14^, hinting at a direct but functionally uncharacterized NTD-SCBD interaction. More recently, we demonstrated that Ca^2+^-induced Sorcin self-association and aggregation can serve as a sensitive functional readout of Sorcin activation^11^. Using stopped-flow light scattering, we showed that Sorcin exhibits relatively low apparent Ca^2+^ sensitivity, becoming activated primarily at Ca^2+^ concentrations near 100 µM^11^. These findings suggested that Sorcin functions as a specialized Ca^2+^ sensor adapted to environments experiencing large transient increases in local Ca^2+^ concentration.

**Figure 1.**
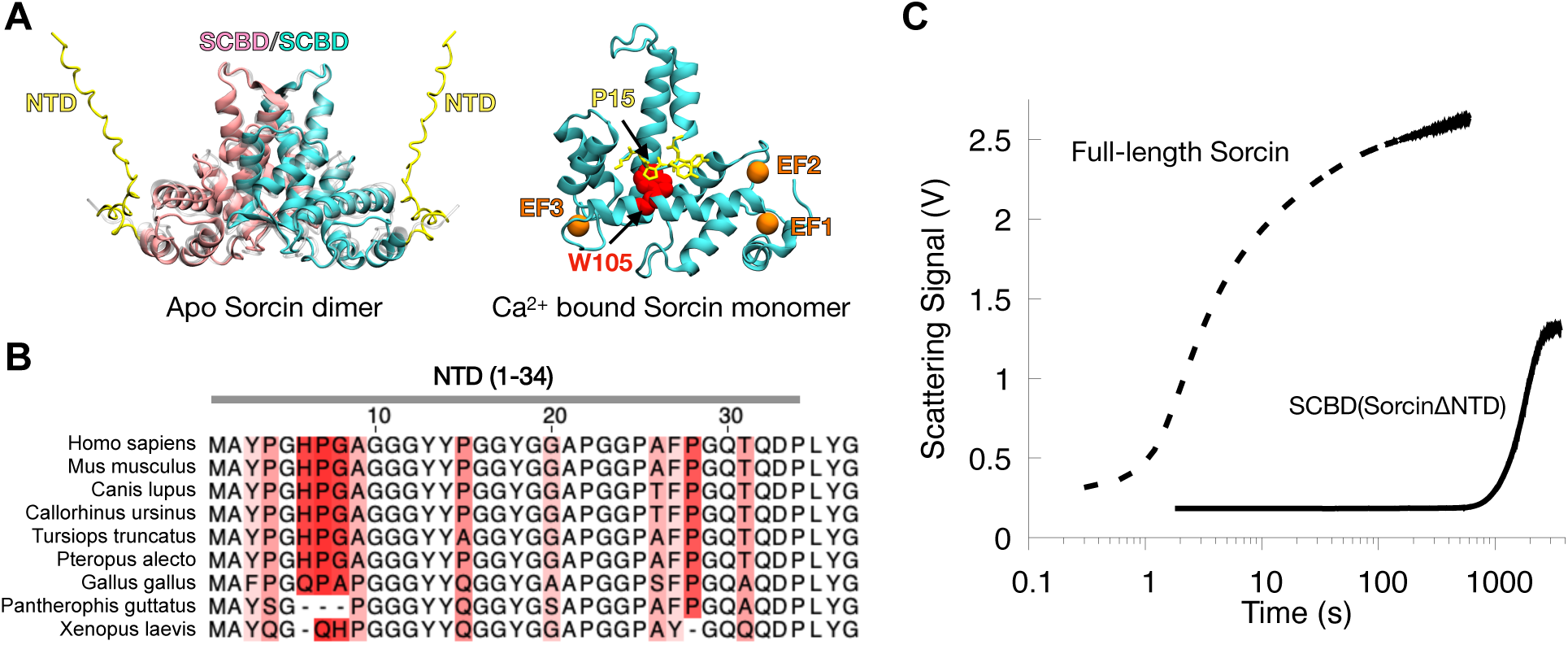
Structural context, sequence conservation, and functional consequences of Sorcin NTD deletion. (A) Left, AlphaFold3-predicted model of the apo Sorcin dimer. The flexible N-terminal domains (NTDs; residues 1-34) are shown in yellow, and the Sorcin calcium-binding domains (SCBDs; residues 35-198) of the two protomers are shown in pink and cyan. The apo crystal structure (PDB 4UPG; resolved for residues 30-198), shown as a transparent overlay, superimposes closely on the AlphaFold3 model. Right: asymmetric unit of the Ca^2+^-bound Sorcin crystal structure (PDB 4USL), with bound Ca^2+^ ions in orange; the contact between Pro15 of the NTD and Trp105 of the SCBD is shown in stick and sphere representation. (B) Multiple sequence alignment of Sorcin from nine representative vertebrate species demonstrates strong sequence conservation within the intrinsically disordered NTD. (C) Stopped-flow light-scattering traces show that deletion of the NTD markedly slows Ca^2+^-induced Sorcin aggregation under matched experimental conditions.

Despite extensive characterization of the SCBD, the function of the NTD remains poorly understood. The NTD is largely absent from available crystal structures because of its conformational flexibility^12,14,28^. Its contribution to Sorcin activation, Ca^2+^ responsiveness, and conformational dynamics has remained unclear. Interestingly, naturally occurring Sorcin isoforms differ primarily in NTD length^13,24^, suggesting that this region may serve a regulatory function. Similar flexible N-terminal extensions are also found throughout the PEF protein family^29–32^, raising the possibility that they represent a conserved mechanism for modulating protein activity and Ca^2+^ responsiveness.

Here we investigated the role of the conserved flexible Sorcin NTD using a combination of biochemical, biophysical, and computational approaches. We generated a truncated construct containing only the calcium-binding domain (SCBD) after NTD deletion and compared its behavior with that of full-length Sorcin. Using Ca^2+^-induced aggregation as a functional readout, we quantified the effects of NTD deletion on Ca^2+^ responsiveness and protein concentration dependence. We further employed multidimensional NMR spectroscopy and long-timescale all-atom MD simulations to elucidate the structural basis of NTD-mediated regulation in both apo and Ca^2+^-bound states. Our results reveal that the NTD acts as a critical regulator of Sorcin activation. The mechanistic studies support a model in which transient, aromatically anchored NTD contacts stabilize activation-associated SCBD surfaces and bias Sorcin toward the Ca^2+^-bound active state. These findings establish an important regulatory role for the NTD in Sorcin function and suggest a broader mechanism by which flexible terminal regions tune the conformational landscapes and signaling properties of EF-hand Ca^2+^ sensors.

## Results

### Sequence conservation suggests an evolutionarily preserved regulatory function of the Sorcin NTD

Sorcin is highly conserved throughout vertebrate evolution^18^. To confirm this, we performed a sequence alignment using nine representative species spanning amphibians, reptiles, birds, and mammals (Figure S1A). Sequence identity between human Sorcin and its mammalian orthologs, including mouse, dog, seal, dolphin, and bat, exceeded 96% for the selected species and isoforms used in the alignment. Remarkably, sequence similarity remained very high even in more distantly related vertebrates, reaching 92.4%, 89.9%, and 89.4% for Sorcin from chicken, snake, and frog, respectively (Table S1). Notably, the disordered NTD of Sorcin also exhibited a high degree of sequence conservation (Figure 1B). This level of conservation suggests that, despite its conformational disorder, the NTD is subject to strong evolutionary constraints, indicating important biological functions that have been maintained throughout vertebrate evolution.

Consistent with the strong conservation observed across species, human Sorcin isoforms and splice variants also exhibit high sequence conservation despite substantial differences in overall length. Most of these length variations result from insertions or deletions within the NTD (Figure S1B). This pattern suggests that the NTD may play a regulatory role, enabling Sorcin to fine-tune its function in different tissues and cell types through the expression of distinct isoforms and splice variants. Furthermore, sequence alignment of all human penta-EF-hand calcium-binding proteins revealed that variation in the length and composition of the glycine-rich NTD is a prominent distinguishing feature within this protein family (Figure S2)^29,32^. Together, these observations suggest that modulation of NTD length and structural properties may represent a conserved regulatory strategy among penta-EF-hand proteins, contributing to their functional diversification and context-dependent regulation.

### Deletion of the N-terminal domain drastically reduces Sorcin responsiveness to Ca^2+^

To further investigate the role of the Sorcin N-terminal domain (NTD) in regulating Sorcin function, we generated a truncated construct lacking the first 34 residues of the NTD, resulting in a protein containing only the Sorcin C-terminal calcium-binding domain (SCBD). We evaluated the response of SCBD to Ca^2+^ using a stopped-flow light-scattering assay previously established in our laboratory to monitor Ca^2+^-induced Sorcin aggregation^11^. Because Sorcin aggregation is triggered by Ca^2+^ binding and the associated conformational changes, this assay provides a sensitive means of assessing the responsiveness and apparent Ca^2+^ sensitivity of Sorcin on the millisecond timescale^11^.

Using this assay, we measured the kinetics of Ca^2+^-induced aggregation of SCBD and compared them with those of full-length Sorcin. Surprisingly, deletion of the glycine-rich intrinsically disordered NTD had a dramatic effect on the protein’s response to Ca^2+^. Under identical experimental conditions, including protein concentration, Ca^2+^ concentration, buffer composition, and temperature, the aggregation lag-time increased from approximately 1 s for full-length Sorcin to nearly 1000 s for SCBD (Figure 1C), indicating that removal of the NTD slows the Ca^2+^-induced aggregation process by approximately three orders of magnitude. Because SCBD aggregated on a minute-to-hour timescale, subsequent Ca^2+^-, protein-, and temperature-dependence measurements were performed by plate-reader based turbidity assay rather than stopped-flow light-scattering assay to increase the throughput of the experiments.

### Ca^2+^ titration reveals reduced Ca^2+^ sensitivity of SCBD

To determine the Ca^2+^ concentration range required for SCBD activation, we performed Ca^2+^ titration experiments using this plate reader-based turbidity assay. Previous stopped-flow measurements of full-length Sorcin showed that aggregation begins at Ca^2+^ concentrations of approximately 60 µM and reaches half-maximal aggregation at 93.7 ± 1.1 µM^11^, a transition likely associated with Ca^2+^ binding to EF3. In contrast, Ca^2+^-induced aggregation of SCBD was not detected until the Ca^2+^ concentration reached approximately 200 µM (Figure 2A). Both the rate and extent of aggregation increased with increasing Ca^2+^ concentration, although the maximal scattering signal approached a plateau above 300 µM Ca^2+^.

**Figure 2.**
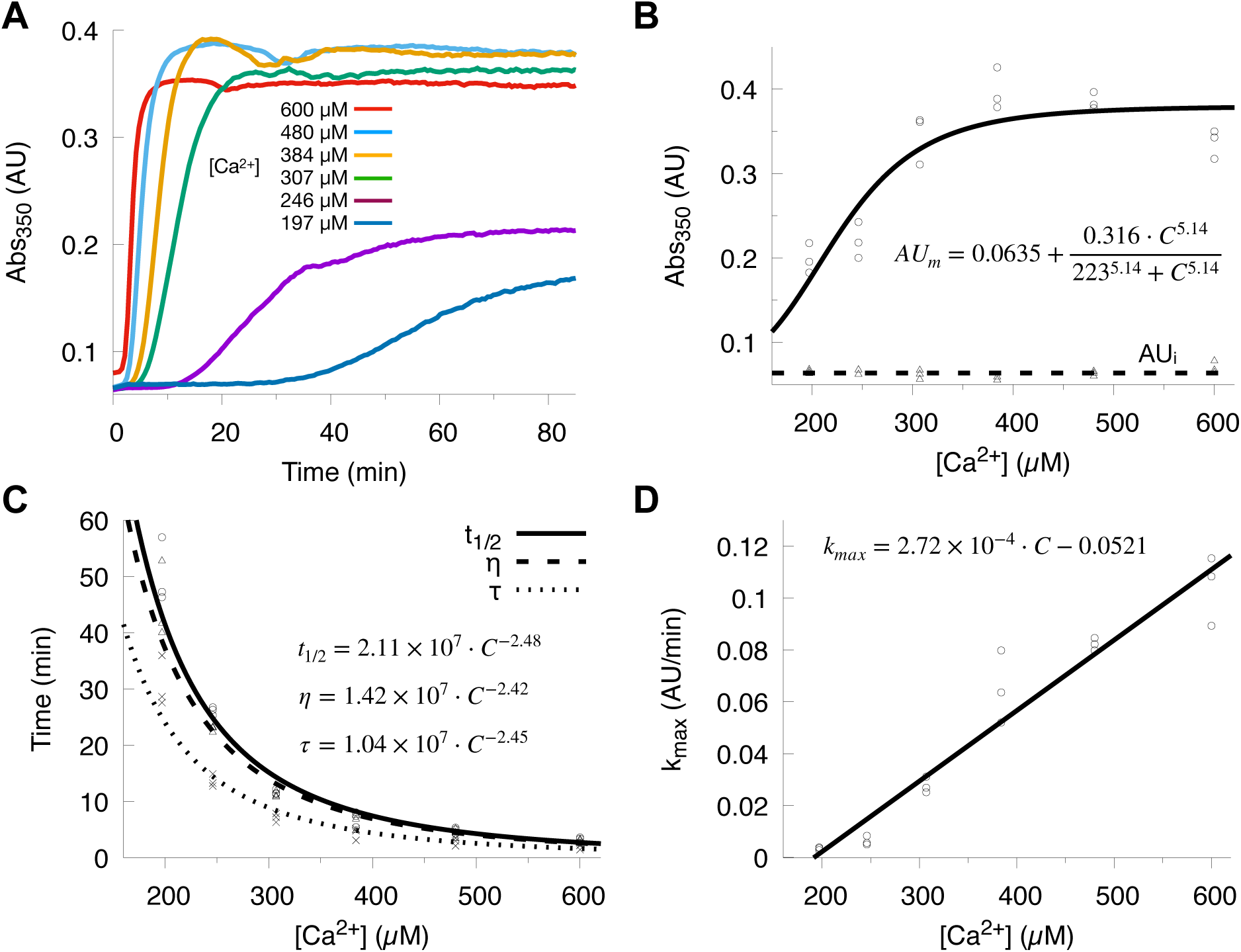
Calcium-concentration dependence of SCBD aggregation kinetics. (A) Representative aggregation time courses for SCBD (10 µM) at Ca^2+^ concentrations ranging from 197 to 600 µM. (B) Dependence of the maximum scattering signal (AUₘ) on Ca^2+^ concentration, fitted with the Hill equation (solid line); the initial signal (AUᵢ) remains essentially constant and is fitted with a horizontal line (dashed line). AUₘ and AUᵢ values from individual experiments are shown as circles and triangles, respectively. (C) The aggregation half-time (t½), the time to the maximum aggregation rate (η), and the lag time (τ) each follow a power law in Ca^2+^ concentration with a scaling exponent of ∼2.5. Individual values of t½, η, and τ are shown as circles, triangles, and crosses, and the power-law fits as solid, dashed, and dash-dotted lines, respectively. (D) The maximum aggregation rate (k_max_) scales linearly with Ca^2+^ concentration (solid line). Individual experiments are shown as circles.

To quantify the aggregation kinetics, we calculated the first and second derivatives of the turbidity curves (Figures S3A and S3B), enabling determination of the maximum aggregation rate (kₘₐₓ) and the corresponding time point (η) at which this rate was achieved. The aggregation kinetics were further analyzed using a double-exponential model as described previously^11^. The fitting yielded key kinetic parameters including the minimum light scattering signal (AU_i_) reflecting the initial background noise, the maximum light scattering signal (AU_m_) reflecting the final extent of aggregation, the aggregation half-time (t_1/2_) at which the signal reaches the midpoint between AU_i_ and AU_m_, and the aggregation lag time (*τ*).

As observed for full-length Sorcin, the dependence of the maximal scattering signal (AU_m_) on Ca^2+^ concentration followed a sigmoidal relationship and was well described by the Hill equation (Figure 2B). Fitting yielded a Hill coefficient of 5.14 ± 2.88 for SCBD, consistent with cooperative Ca^2+^-induced aggregation, although the large uncertainty precludes a precise comparison with the value of 9.15 ± 0.89 measured for full-length Sorcin^11^. More strikingly, the half-maximal aggregation concentration increased to 223 ± 20.6 µM, more than twofold higher than that of full-length Sorcin (93.7 ± 1.1 µM). These results demonstrate that deletion of the N-terminal domain markedly reduces the apparent Ca^2+^ sensitivity of Sorcin.

Analysis of the kinetic parameters further revealed that all characteristic times, including the aggregation half-time (t_1/2_), the lag time (*τ*), and the time required to reach the maximum aggregation rate (η), followed a power law with a negative scaling exponent of about -2.5 as Ca^2+^ concentration increased (Figure 2C), indicating that elevated Ca^2+^ strongly accelerates the aggregation process. In contrast, the maximum aggregation rate (kₘₐₓ) increased approximately linearly with Ca^2+^ concentration across the range examined (Figure 2D).

### Deletion of the NTD increases the protein concentration threshold for aggregation

In addition to Ca^2+^ concentration, protein concentration is another key parameter that influences Sorcin aggregation behavior^11^. To investigate how NTD deletion affects the sensitivity of aggregation to protein concentration, we performed SCBD concentration titration experiments using the plate reader-based turbidity assay. As expected, increasing SCBD concentration accelerated the aggregation kinetics and resulted in a greater extent of aggregation (Figure 3A). The corresponding first and second derivatives of the turbidity curves are shown in Figures S4A and S4B. The maximal scattering signal (AU_m_) increased linearly with SCBD concentration (Figure 3B). Similar to the Ca^2+^ concentration dependence, the characteristic times (t_1/2,_ η, and *τ*) followed a power law with a negative scaling exponent of about -3 as SCBD concentration increased, whereas kₘₐₓ increased linearly with SCBD concentration (Figures S4C and S4D).

**Figure 3.**
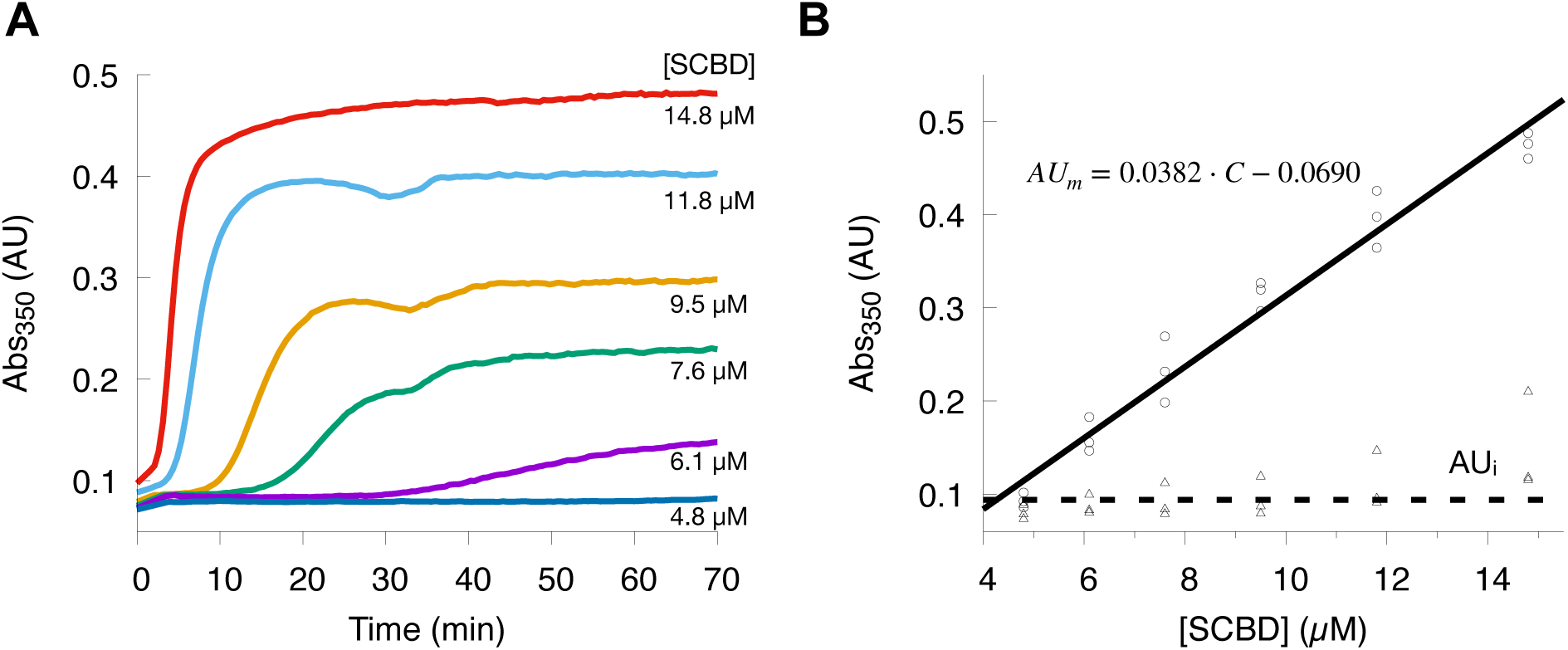
Protein-concentration dependence of SCBD aggregation kinetics. (A) Representative aggregation time courses for SCBD at concentrations ranging from 4.8 to 14.8 µM in the presence of 300 µM Ca^2+^. (B) Dependence of the maximum scattering signal (AUₘ) on SCBD concentration, fitted with a linear equation (solid line); the initial signal (AUᵢ) remains essentially constant and is fitted with a horizontal line (dashed line). AUₘ and AUᵢ values from individual experiments are shown as circles and triangles, respectively.

Although aggregation of both full-length Sorcin and SCBD is strongly dependent on protein concentration, the minimum concentration required to observe measurable aggregation differs substantially between the two constructs. For full-length Sorcin, aggregation could be detected at concentrations as low as 0.4 µM^11^. In contrast, SCBD required concentrations of at least 5 µM to exhibit detectable aggregation, representing more than a tenfold increase in the concentration threshold. These results indicate that SCBD is significantly less prone to self-association and aggregation, requiring not only higher Ca^2+^ concentrations but also substantially higher protein concentrations to initiate aggregation. Together with the Ca^2+^ titration results, these findings demonstrate that deletion of the NTD markedly attenuates Sorcin responsiveness, resulting in a substantially more inert Ca^2+^ sensor.

### Arrhenius analysis for the temperature dependence of SCBD aggregation

The adaptation of the plate reader-based turbidity assay also allowed us to investigate how SCBD aggregation depends on temperature. Such characterization was not possible for full-length Sorcin because our stopped-flow instrument lacks a temperature-control unit. As shown in Figure 4A, increasing temperature markedly accelerated SCBD aggregation kinetics without altering the plateau turbidity signal, indicating that temperature affects the rate of aggregation but not the total extent of aggregation achieved at equilibrium.

**Figure 4.**
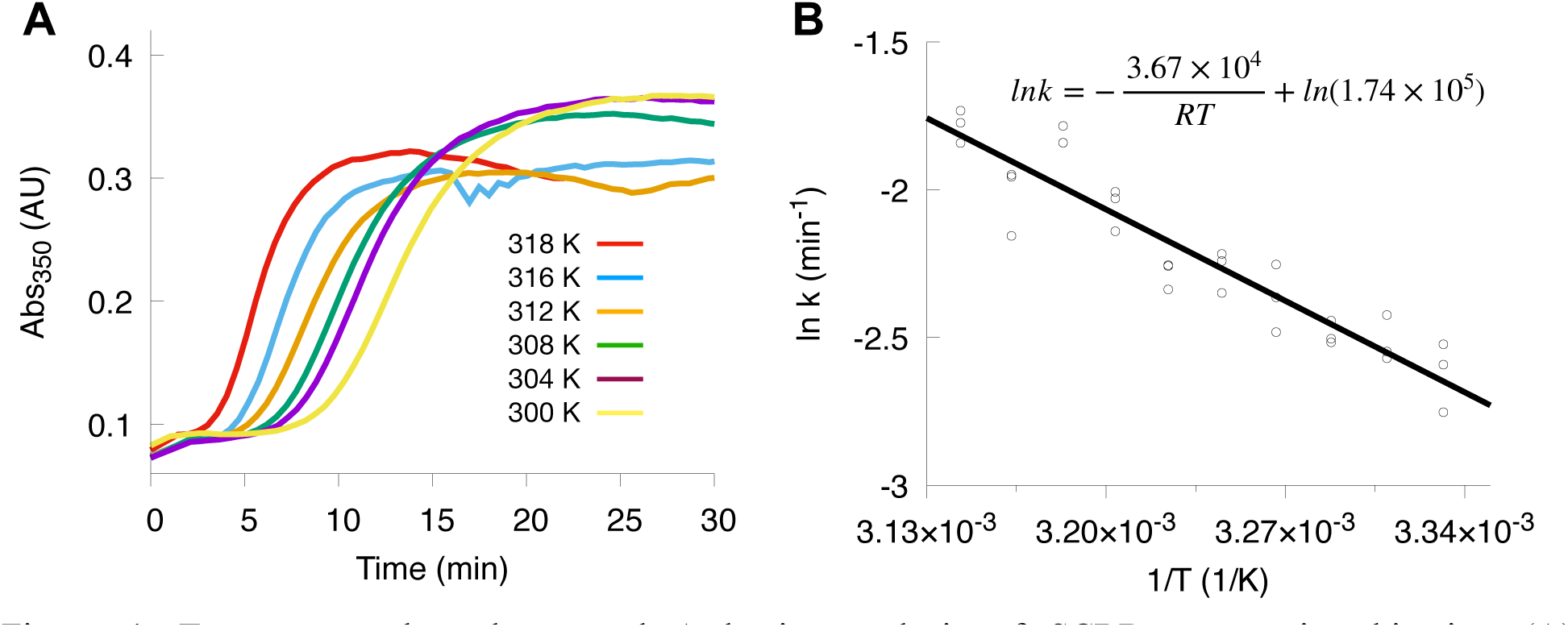
Temperature dependence and Arrhenius analysis of SCBD aggregation kinetics. (A) Representative aggregation time courses for SCBD in the presence of 300 µM Ca^2+^ at temperatures ranging from 300 to 318 K. (B) The aggregation rate, characterized by the inverse of the half-time (t_1/2_), follows Arrhenius behavior, yielding an apparent activation energy of 36.7 kJ/mol. Values of 1/t_1/2_ are shown as circles and the Arrhenius fit as a solid line.

Arrhenius analysis of the aggregation half-times yielded an apparent activation energy of 36.7 kJ·mol^-1^ (Figure 4B), a relatively modest value for a protein self-association process. This apparent activation energy reflects the temperature dependence of the overall aggregation pathway and likely encompasses multiple coupled events, including Ca^2+^ binding, Ca^2+^-induced conformational rearrangements, and nucleation processes that precede aggregate growth. Because turbidity measurements report the macroscopic formation of light-scattering particles rather than individual molecular events, the derived value should be interpreted as an apparent activation energy for the overall aggregation process rather than as the microscopic activation barrier of any single elementary step. The moderate barrier is consistent with aggregation following Ca^2+^-induced exposure of self-association surfaces rather than requiring global protein unfolding^11,14^.

### NTD deletion perturbs the EF-hand calcium-binding core of the SCBD

To understand the molecular mechanism by which the N-terminal domain (NTD) regulates Sorcin conformation and dynamics, we performed three-dimensional heteronuclear NMR experiments on SCBD using purified ^2^H/^15^N/^13^C triple-labeled protein in the absence of Ca^2+^. The two-dimensional ^1^H-^15^N HSQC spectrum displayed well-dispersed resonances, indicating that SCBD is well folded and structurally stable (Figure 5A). The overall spectrum closely resembles that of full-length Sorcin reported previously by us^27^, suggesting that deletion of the NTD does not alter the global fold of the calcium-binding domain.

**Figure 5.**
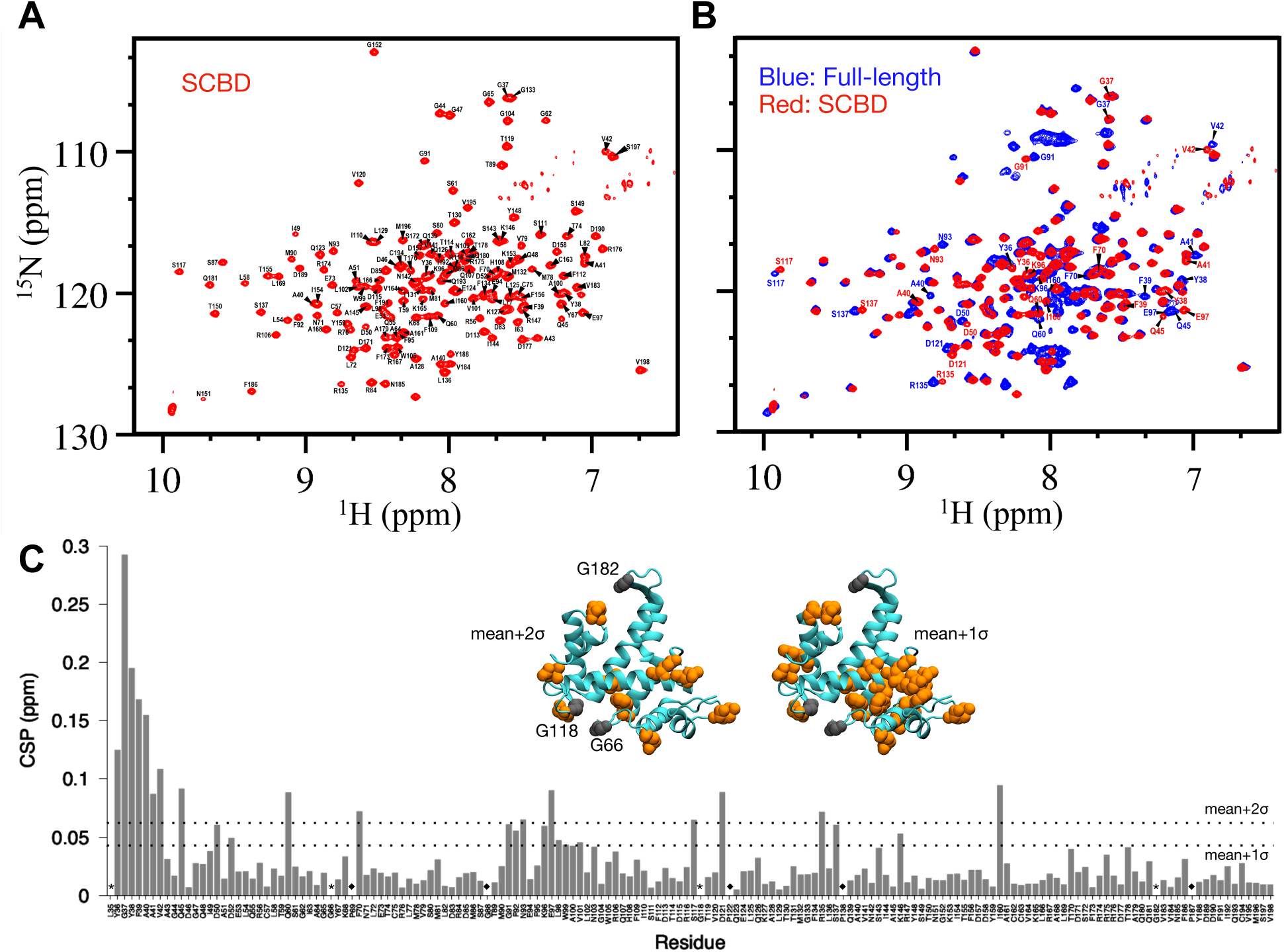
Chemical shift perturbations in SCBD upon NTD deletion. (A) Backbone assignment and annotation of the ¹H-15N HSQC spectrum of triple-labeled SCBD. (B) Overlay of the ¹H-¹⁵N HSQC spectra of SCBD (red) and full-length Sorcin (blue); residues with large chemical shift perturbations (CSPs) are labeled. (C) Per-residue CSP between SCBD and full-length Sorcin. Dashed lines indicate the mean + 1σ and mean + 2σ thresholds. The first eight residues were excluded from the analysis owing to their proximity to the NTD. Residues exceeding the mean + 1σ and mean + 2σ thresholds are mapped onto the apo Sorcin crystal structure (PDB 4UPG) in the inset (orange). Residues whose peaks disappeared upon NTD deletion are marked with asterisks and mapped onto the structure in gray; residues absent from both the SCBD and full-length spectra are marked with a diamond.

To investigate the structural and dynamic effects of NTD deletion in greater detail, we independently completed backbone resonance assignments for SCBD (Figure 5A) and compared the resulting ^1^H, ^15^N, and ^13^C chemical shifts with those of full-length Sorcin (Figure 5B)^27^. The residue-specific chemical shift perturbations (CSPs), calculated from the backbone amide ¹H and ¹⁵N shifts, are shown in Figure 5C. If the NTD did not interact with the SCBD, minimal CSPs would be expected across the protein. In contrast, we observed measurable perturbations at multiple sites within the SCBD. Although most were small in magnitude, a subset of residues displayed larger changes. Residues 35-42 were excluded from the analysis because their shifts were likely dominated by the new N-terminus. For the remaining residues, the mean CSPs was 0.024 ± 0.019 ppm. Nineteen residues exceeded the descriptive mean + 1 standard deviation (σ) threshold (>0.043 ppm): Gln45, Asp50, Asp52, Gln60, Phe70, Gly91, Phe92, Asn93, Lys96, Glu97, Leu98, Trp99, Val101, Ser117, Asp121, Arg135, Ser137, Lys146, and Ile160. Nine exceeded mean + 2σ (>0.062 ppm): Gln45, Gln60, Phe70, Asn93, Glu97, Ser117, Asp121, Arg135, and Ile160.

Rather than distributing randomly across the domain, these residues clustered within the calcium-sensing core and its associated structural elements (Figure 5C inset). Four spatial clusters were apparent. The first comprised the EF1 calcium-binding loop and its flanking regions (Gln45, Asp50, Asp52, Gln60, and Phe70), including two of the acidic residues that coordinate Ca^2+^ in EF1. The second, and most contiguous, extended from the EF2 loop through the D-helix (Gly91, Phe92, Asn93, Lys96, Glu97, Leu98, Trp99, and Val101). Because the D-helix is the pivot element that undergoes the large reorientation accompanying Ca^2+^-induced activation and this cluster includes aromatic and hydrophobic residues that contribute to the activation-exposed surface^14^, its perturbation is of particular mechanistic interest. The third cluster fell within the EF3 calcium-binding loop (Ser117 and Asp121), with Ser117 itself a Ca^2+^-coordinating residue. The fourth mapped to the EF3-EF4 junction and the dimerization subdomain (Arg135, Ser137, Lys146, and Ile160 on the G-helix). Collectively, these residues delineate the EF1-EF3 calcium-binding loops, the D-helix, and the adjoining surfaces that reorganize upon activation. We adopted the mean-plus-1σ set for subsequent comparison with the MD simulation data, as this more inclusive list preserves the full spatial extent of each cluster. The mean-plus-2σ subset marks the core of each cluster.

In addition, three assigned full-length resonances (Gly66, Gly118, and Gly182) were absent from the SCBD spectrum. All three are glycines located in EF-hand loop or inter-EF turn regions: Gly118 is the conserved glycine within the EF3 calcium-binding loop, whereas Gly66 and Gly182 occupy loop/turn positions in the EF1-EF2 and EF5 regions, respectively. The loss of these resonances indicates that, following NTD removal, the corresponding backbone amides are broadened beyond detection, most plausibly by intermediate-timescale conformational exchange between alternative loop geometries or by faster solvent exchange upon loss of NTD-mediated shielding. Because these residues were readily observed in full-length Sorcin, their disappearance in SCBD reflects a genuine, localized increase in dynamics at the EF-hand loops rather than an intrinsic property of these glycines.

Taken together, the residues with significant chemical shift perturbations and those with disappearing resonances converge on the same region of the protein: the EF-hand calcium-binding loops (EF1-EF3), the D-helix, and the surrounding elements that constitute the Ca^2+^-sensing and target-recognition surfaces of the SCBD (Figure 5C inset). This convergence indicates that the structural consequences of NTD deletion are focal rather than global and are centered on the calcium-binding core.

### Comparison of NMR perturbations and MD contact analysis distinguishes direct NTD-SCBD contacts from allosteric effects

While the NMR analysis identified the SCBD residues whose environment is altered by NTD deletion, CSPs cannot by themselves distinguish residues that are directly contacted by the NTD from those affected by allosteric propagation of a conformational change. Moreover, because Sorcin is a homodimer, the perturbations cannot indicate whether a given change arises from an intramolecular contact within a protomer or an intermolecular contact from the partner protomer. To resolve these ambiguities, we reanalyzed our previously reported 10-µs MD simulation of apo Sorcin in the Ca^2+^-free inactive state^33^, the same state probed by NMR here, to obtain a residue-resolved map of NTD-SCBD contacts.

For each SCBD residue we calculated an occupancy orientated from NTD contact, defined as the fraction of trajectory frames in which the residue lay within 4.5 Å (heavy atom to heavy atom) of any NTD residue. Because CSPs report a single, protomer-averaged environment for each residue and cannot distinguish which NTD is responsible for a contact, we combined all NTD contacts (intramolecular and intermolecular from both protomers) into a single per-residue occupancy for comparison with the NMR data (Figure 6A). Similar to the CSP analysis, the N-terminal helix of the SCBD (residues 35-42) was excluded because of its covalent adjacency to the NTD. Rather than binding a single defined site, the NTD sampled a broad but non-random set of SCBD surfaces through a network of transient contacts, with occupancy concentrated in three contiguous regions that together form one face of the calcium-binding subdomain: the C-terminal segment of EF1 and the EF1-EF2 linker (residues ∼59-69), the D-helix (residues ∼92-110), and the entrance to the EF3 calcium-binding loop (residues ∼116-117). No single contact approached permanence, indicating an intrinsically dynamic, multimodal interaction rather than a fixed association. This analysis is insensitive to the cutoff (4.5 Å) chosen to define contact. The results with cutoff 4.0 or 5.0 Å were shown in Figure S5.

**Figure 6.**
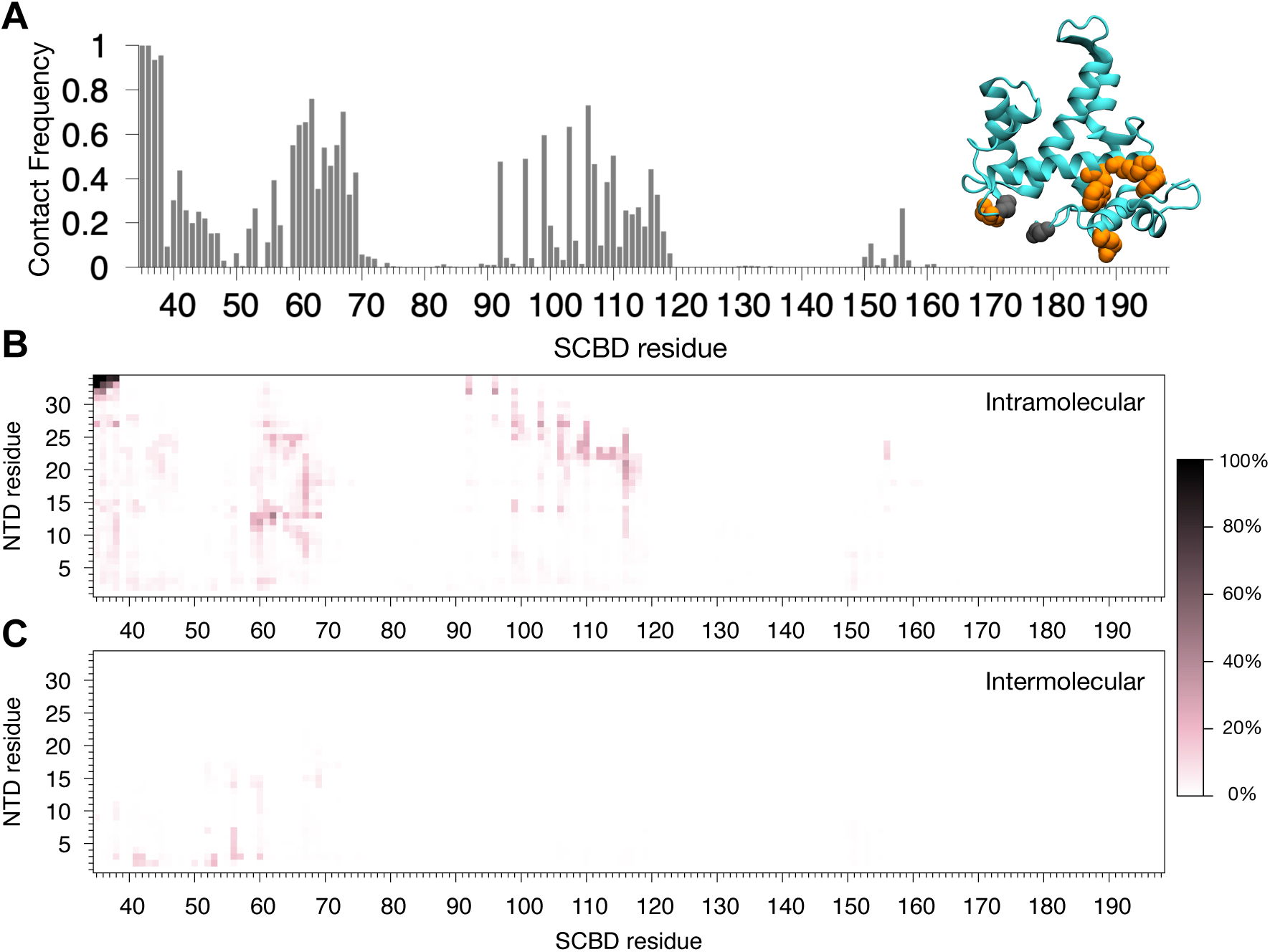
Molecular dynamics simulations reveal the interaction between the Sorcin NTD and SCBD in the apo state. (A) Per-residue contact frequency of the SCBD with the NTD (1 denotes 100% occupancy), defined as the fraction of frames in which any NTD heavy atom lies within 4.5 Å of a given SCBD residue. Residues that are both highly occupied in the simulation and strongly perturbed in the NMR analysis are mapped onto the apo crystal structure (inset, orange). Gly66 and Gly118, whose resonances disappear upon NTD deletion in the NMR spectra, also contact the NTD in these simulations. (B) Heat map of the contact frequency between NTD and SCBD residues arising from intramolecular (within-protomer) interactions. (C) Heat map of the contact frequency between NTD and SCBD residues arising from intermolecular (between-protomer) interactions.

Several CSP sites coincided with substantial occupancy, including Gln60 (0.68), Phe92 (0.48), Lys96 (0.49), Trp99 (0.60), and Ser117 (0.32), supporting possible direct effects (Figure 6A inset). Other strongly perturbed sites had little contact: Glu97 (0.04), Asp121, Arg135, and Ile160 (≤0.01). Their perturbations are therefore more consistent with locally propagated or distal allosteric responses. Among the three disappearing resonances, Gly66 and Gly118 occurred in contacted regions and Gly182 had no detected contact (Figure 6A inset). This comparison supports both direct and distal consequences of NTD deletion.

To identify which NTD residues are responsible for these contacts, we examined how NTD in each protomer interacts with SCBDs and produced intramolecular and intermolecular residue-residue contact maps shown in Figures 6B and 6C. Both were dominated by transient interactions, with no residue pair exceeding a contact probability of 45% after excluding NTD-SCBD junction residues, consistent with a dynamic, multivalent interface. Intramolecular contacts were the more prevalent of the two and were mediated predominantly by the aromatic-rich central segment of the NTD (residues ∼12-27), in particular Tyr13, Tyr14, Tyr18, and Phe27, which engaged precisely the surfaces identified above: the EF1-EF2 linker, the D-helix, and the EF3-loop entrance. Tyr13 and Tyr14 belong to the conserved 12-GYYPGG-17 motif whose Pro15 contacts Trp105 in the Ca^2+^-bound crystal structure^14^. Thus, apo contacts involving this aromatic motif may sample surfaces that become reorganized during activation. Intermolecular contacts were weaker (maximum probability ∼19%) and originated from a distinct region, the extreme N-terminal residues 2-7 (most notably Tyr3), which reached across the dimer interface to the EF1 region (residues 41-60) of the partner protomer. Thus, the NTD predominantly folds back onto the calcium-binding domain of its own protomer, engaging it through its conserved aromatic residues, while its N-terminal tip can additionally bridge to the neighboring protomer.

### The NTD stabilizes the Ca^2+^-bound active state through a dynamic ensemble of multimodal contacts

The combination of NMR spectroscopy and MD simulations provided detailed insight into the effects of the N-terminal domain (NTD) on Sorcin structure and dynamics in the absence of Ca^2+^. However, Ca^2+^-induced self-association and aggregation prevented NMR characterization of the NTD’s role in the Ca^2+^-bound state. We therefore used long-timescale all-atom MD simulations in explicit solvent on Anton 3^34^ to directly investigate the dynamics of Sorcin with and without the NTD in the Ca^2+^-bound state. Previously, we reported four 10-µs MD simulations of full-length Sorcin in the Ca^2+^-bound state^33^, both with and without distance restraints applied between the bound Ca^2+^ ions and EF hands. Here we performed four corresponding 10-µs simulations of the isolated calcium-binding domain (SCBD) following NTD deletion (Movie S1-4), using identical protocols to enable a direct, matched comparison with the full-length protein^33^. A summary of the four simulations performed for this study and the previously generated simulations are given in Table S2.

In unrestrained SCBD, Ca^2+^ rapidly dissociated from all three EF hands (Figure S6), as observed previously for full-length Sorcin. Three additional simulations retained Ca^2+^ using minimal weak, extensive weak, or extensive strong coordination restraints as described in the Method secion and defined previously^33^, allowing matched comparison of the constructs. The C_α_-RMSD of each construct relative to the crystallographic Ca^2+^-bound active structure (residues 35-198) is shown in Figure 7A-D. All simulations diverged from the starting active conformation to varying degrees, even when Ca^2+^-coordination restraints were applied.

**Figure 7.**
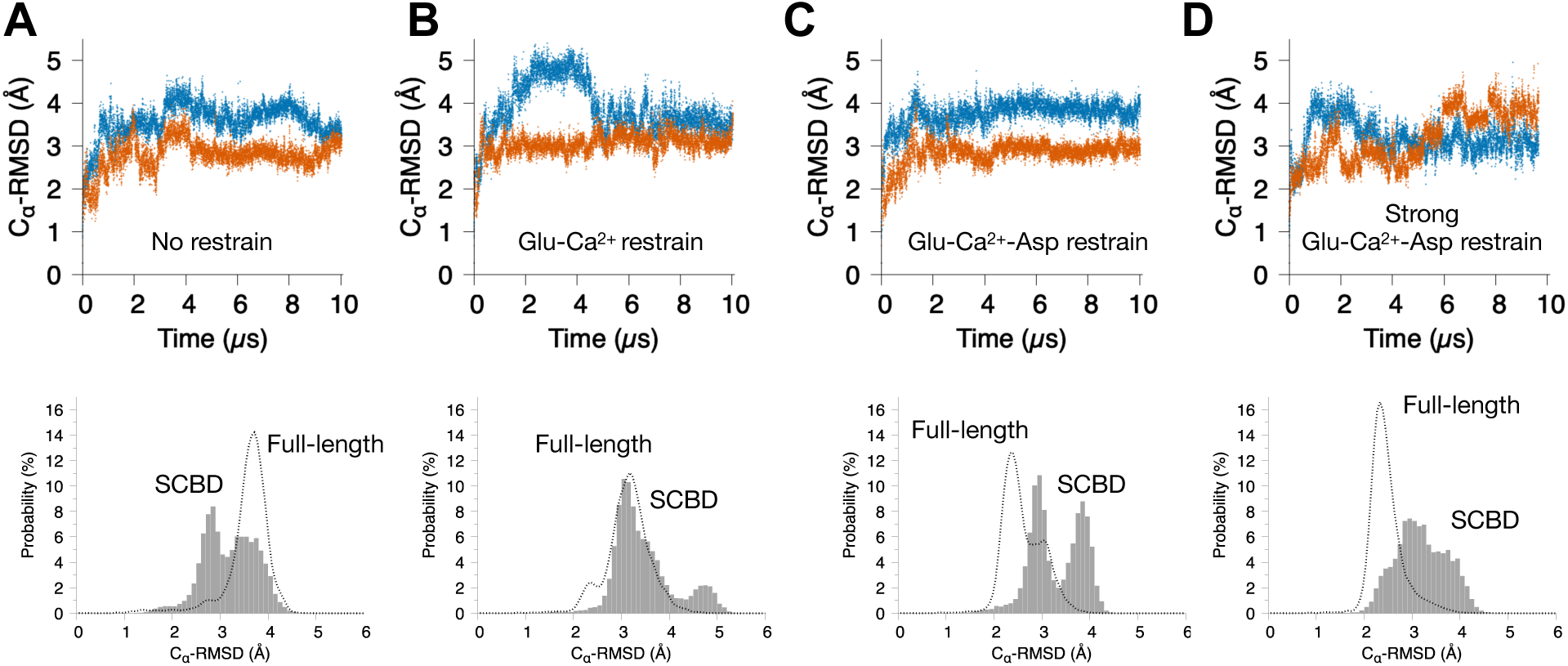
Structural stability and time evolution of SCBD initiated from the Ca^2+^-bound active state (PDB 4USL). (A) C_α_-RMSD of chains A and B of SCBD with no distance restraints between Ca^2+^ and the EF-hands. (B) C_α_-RMSD with weak distance restraints between Ca^2+^ and the bidentate glutamate of each EF-hand. (C) C_α_-RMSD of chains A and B of SCBD with weak distance restraints between Ca^2+^ and all charged coordinating residues of each EF-hand. (D) C_α_-RMSD of chains A and B of SCBD with strong distance restraints between Ca^2+^ and all charged coordinating residues of each EF-hand. The Ca^2+^-bound Sorcin structure (PDB 4USL) was used as the reference for alignment and RMSD calculation in all trajectories. Beneath each time trace, the C_α_-RMSD distribution from the corresponding SCBD simulation is shown and compared with that of full-length Sorcin under the same restraint scheme.

The key comparison is how full-length Sorcin and SCBD responded to increasing restraint strength. As the restraints held Ca^2+^ more effectively within the EF-hands, full-length Sorcin progressively converged toward the active state, its mean C_α_-RMSD decreasing from ∼3.3 Å under minimal restraints to ∼2.4 Å under the strongest restraints (Figures 7B and 7D, bottom portion). SCBD, in contrast, failed to converge and remained substantially more divergent (∼3.4-3.5 Å) across all restrained conditions (Figure 7, bottom portion). Consequently, the difference between the two constructs grew monotonically with restraint strength, from negligible under minimal restraints to ∼1.1 Å under the strongest restraints, where full-length Sorcin was well-stabilized in the active conformation, but SCBD was not. The distribution also became broader for SCBD compared to full-length Sorcin (Figures 7D, bottom portion). The divergence was frequently asymmetric between the two protomers, with one chain of SCBD deviating markedly more than the other (Figure 7A-D), consistent with the conformational asymmetry of the Sorcin dimer observed in our earlier simulations^33^. In the unrestrained simulations, both constructs diverged and Ca^2+^ dissociated, precluding a meaningful comparison in that condition. Together, these results indicate that full-length Sorcin can be held in the crystallographic active conformation when its EF-hand Ca^2+^ coordination is maintained, whereas the isolated SCBD cannot, demonstrating that the NTD contributes directly to stabilizing the Ca^2+^-bound active conformation.

To identify the molecular basis of this stabilization, we reanalyzed the full-length Sorcin trajectory in which Ca^2+^ ions were restrained within the EF-hand binding sites. We focused on the simulation employing extensive, strong restraints, as the RMSD analysis established that this condition best maintains the crystallographic active state (Figures 7D, bottom portion). We computed intramolecular and intermolecular NTD-SCBD contact maps for this trajectory, excluding the covalently connected residues at the NTD-SCBD junction (Figure 8A and 8B). This analysis confirmed the crystallographic observation that NTD folds back and inserts into a hydrophobic pocket of the SCBD, contacting Trp105 through Pro15 of the conserved GYYPGG motif (Figure 8C-E)^14^. This Pro15-Trp105 interaction occurred with a contact probability of 0.285 and ranked among the ten most frequent NTD-SCBD contacts in our simulation, providing a mutual validation between the crystallographic structure and the simulations^14^. Interestingly, in apo Sorcin the NTD frequently sampled the D-helix near Trp105 but rarely contacted Trp105 itself (Figure 6A). In contrast, exposure of the active-state hydrophobic pocket in the Ca^2+^-bound Sorcin enabled recurrent Pro15 docking. Trp105 was additionally engaged by Tyr14 (probability 0.263), indicating that the pocket is contacted by more than one NTD aromatic residue.

**Figure 8.**
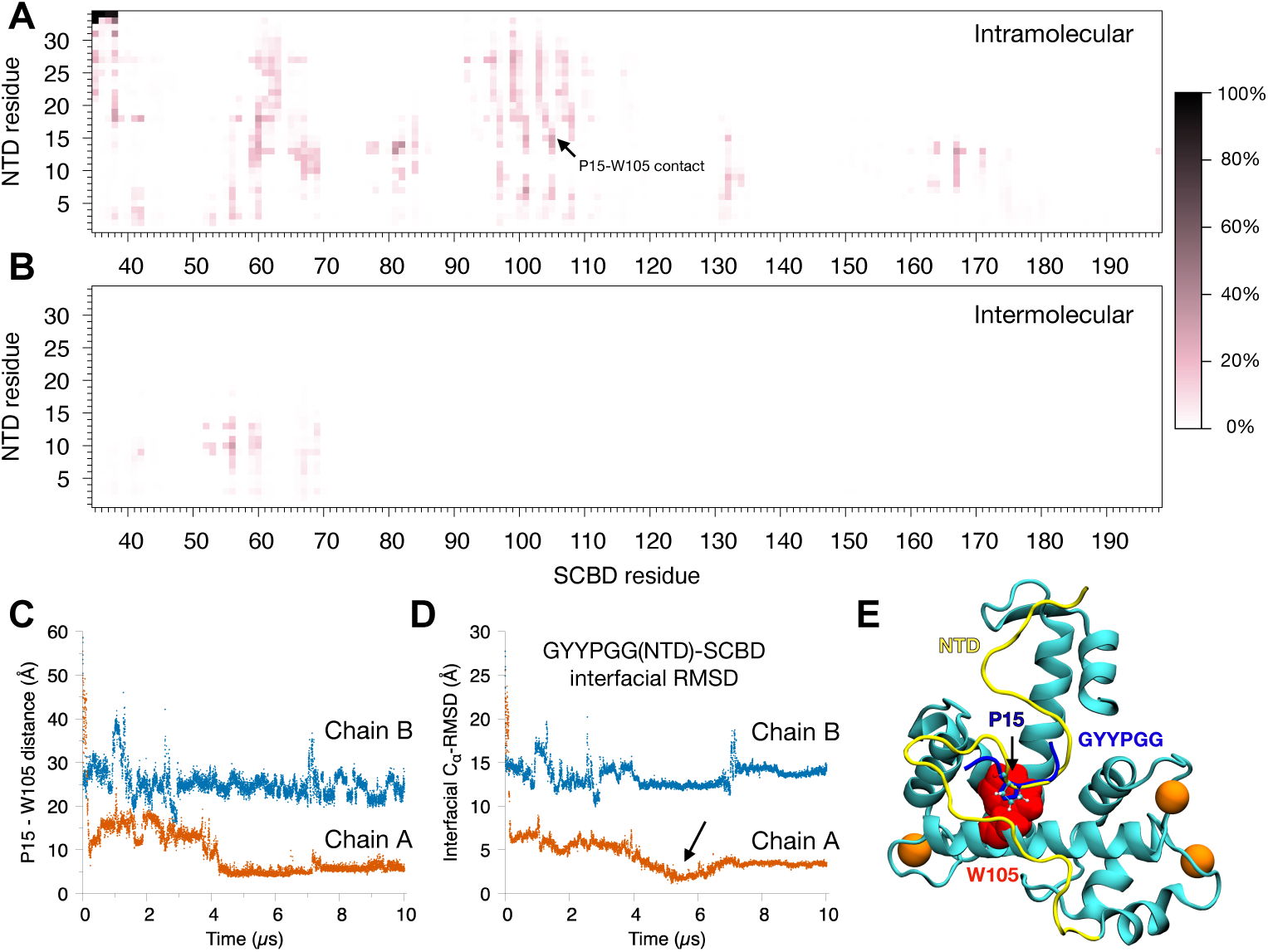
Molecular dynamics simulations reveal the interaction between the Sorcin NTD and SCBD in the Ca^2+^-bound active state. Strong distance restraints were applied between Ca^2+^ and all charged coordinating residues of each EF-hand to maintain the full-length dimer in the active state. (A) Heat map of the contact frequency between NTD and SCBD residues arising from intramolecular (within-protomer) interactions. (B) Heat map of the contact frequency between NTD and SCBD residues arising from intermolecular (between-protomer) interactions. (C) Distance between Pro15 of the NTD and Trp105 of the SCBD over time; a close contact is established and maintained after ∼4 µs in chain A. (D) Time course of the interfacial RMSD between the NTD GYYPGG motif and its crystallographically resolved binding pose on the SCBD; a docked configuration closely matching the crystal structure is reached at 5.16 µs (arrow), with an interfacial RMSD of 1.2 Å. (E) Snapshot of full-length Sorcin (chain A) at t = 5.16 µs (NTD, yellow; SCBD, cyan; Trp105, red), resembling the Ca^2+^-bound crystallographic state; the crystallographic pose of the ¹²GYYPGG¹⁷ motif is shown in blue for comparison.

The analysis further revealed that the Pro15-Trp105 interaction is not the dominant contact. As shown in Figure 8A, several other interactions occurred with equal or greater frequency, including contacts between the GYYPGG aromatics Tyr13 and Tyr14 and the EF2 calcium-binding loop (residues 81-82; probabilities up to 37%), between residue Pro7 and the D-helix (residue 101; 32%), between Tyr18 and the EF1-EF2 linker (residue 60; 32%), and between Tyr13 and the EF5 region (residue 167; 31%). Intermolecular contacts of comparable magnitude were also observed, in which the N-terminal segment of the NTD (residues 9-13) engaged the EF1 region of the partner protomer (residues 55-60; probability up to 31%). As in the apo state, these interactions were mediated predominantly by the conserved aromatic residues of the NTD and were individually transient, with no single contact approaching permanence. The crystallographic contact of Pro15-Trp105 therefore represents one mode within a broader, aromatically anchored network spanning the EF2 loop, D-helix, EF1 linker, and EF5 region. This multimodal network provides a structural basis for NTD-dependent stabilization of the active ensemble.

## Discussion

Sorcin is a Ca^2+^-binding protein that functions as a cellular Ca^2+^ sensor^11,14,35^. In our previous work, we established that Ca^2+^-induced Sorcin aggregation can serve as a convenient functional readout of Sorcin activation and Ca^2+^ responsiveness^11^. Using this approach, we discovered that Sorcin exhibits relatively low Ca^2+^ sensitivity, with activation occurring primarily in the ∼100 µM Ca^2+^ range^11^. Subsequently, using long-timescale all-atom MD simulations on the Anton 3 supercomputer, we proposed a coordination-biased conformational selection mechanism for Sorcin activation^33^. In the present study, we identify the N-terminal domain (NTD) as a critical structural element that regulates Sorcin activation and its responsiveness to Ca^2+^ binding. Deletion of the NTD, generating the Sorcin calcium-binding domain construct (SCBD), substantially reduces the protein’s responsiveness to Ca^2+^. SCBD requires approximately twofold higher Ca^2+^ concentrations and more than tenfold higher protein concentrations to initiate detectable aggregation, a process that reflects Ca^2+^-induced conformational activation.

To uncover the molecular basis of this regulation, we combined NMR spectroscopy and long-timescale MD simulations on Anton 3. In the apo state, NMR chemical shift perturbations, interpreted together with an MD contact analysis, indicate that the NTD makes direct but transient contacts with the calcium-binding domain while also exerting allosteric effects. Notably, these perturbations are focused on the EF-hand calcium-binding loops and the adjoining D-helix rather than distributed uniformly across the domain. Complementary all-atom MD simulations of the Ca^2+^-bound state revealed that the NTD engages the calcium-binding domain through a dynamic ensemble of multimodal interactions that stabilize the active conformation observed in crystal structures, with the previously crystallized NTD-GYYPGG-motif and CTD-hydrophobic-pocket contact representing only one of many recurrent binding modes^14^. Together, these findings support a model in which the NTD promotes and stabilizes the Ca^2+^-bound, aggregation-prone active state of Sorcin. Consequently, NTD deletion shifts the conformational equilibrium away from this active state, thereby reducing the propensity for Ca^2+^-induced self-association and aggregation.

Although our data strongly support this NTD-mediated stabilization model, alternative mechanisms cannot be completely excluded. One possibility is that the NTD participates directly in intermolecular interactions during aggregation. For example, NTDs from one Sorcin dimer may interact with the calcium-binding domain of another dimer, or NTD-NTD interactions may contribute to nucleation and aggregate growth through multivalent intermolecular contacts. Consistent with this possibility, our contact analysis detected intermolecular NTD-SCBD contacts within the dimer, in which the N-terminal segment of one protomer reached across the interface to the calcium-binding domain of the other. Distinguishing among these possibilities will require structural characterization of aggregation intermediates and aggregated states, which remain challenging to study using either solution NMR or atomistic MD simulations. However, our data argue against a model in which the NTD functions primarily by modulating Ca^2+^ binding affinity. Previous simulations demonstrated that Ca^2+^ association and dissociation in full-length Sorcin are both extremely rapid and near the diffusion limit. Deletion of the NTD does not substantially alter these kinetic properties, and Ca^2+^ remains weakly bound in SCBD. Furthermore, changes in Ca^2+^ affinity alone are unlikely to account for the approximately three orders of magnitude reduction in aggregation rate observed upon NTD deletion.

In addition, the simulations also do not establish converged thermodynamic populations. Contact occupancies and RMSD distributions should therefore be interpreted as semi-quantitative tendencies. The physical accessibility of NTD-SCBD contacts and the greater departure of matched SCBD trajectories from the active structure are mechanistically informative, but independent and much longer replicate simulations will be needed for rigorous population estimates.

One of the most surprising findings from our previous studies was that Sorcin responds to Ca^2+^ concentrations in the ∼100 µM range^11^, considerably higher than those required to activate many other EF-hand Ca^2+^ sensors^7,9^. Such concentrations are typically reached only transiently within specialized cellular microdomains or highly excitable tissues, such as cardiomyocytes and skeletal muscle cells^36,37^. The further reduction in Ca^2+^ sensitivity observed for SCBD, with activation shifted toward the ∼200 µM range, suggests that NTD truncation may restrict Sorcin activity to environments experiencing exceptionally high local Ca^2+^ concentrations. An intriguing example is the recently discovered mitochondrial isoform of Sorcin, which has an estimated molecular weight of 18.5kDa^24^. Its mitochondrial location does demand a tolerance to high Ca^2+^ concentration and low Ca^2+^ sensitivity. In fact, naturally occurring Sorcin isoforms and splice variants differ primarily in the length of their NTDs. This observation raises the possibility that evolutionary modulation of NTD length provides a mechanism for tuning Sorcin responsiveness to distinct cellular Ca^2+^ signaling environments. Future studies examining the tissue-specific expression patterns and functional properties of these isoforms may provide important insights into this regulatory strategy.

The regulatory role of the NTD may extend beyond Sorcin. Glycine-rich N-terminal extensions are common across the penta-EF-hand family and are a major source of its structural diversity^14,29–32^, suggesting that flexible NTDs may serve analogous regulatory functions throughout the family. Similar N-terminal regulation has been described for other EF-hand proteins, including calcineurin B homologous protein 3 (CHP3) and recoverin^38–40^, indicating that flexible terminal regions may be a general strategy for modulating the conformational landscapes and activation properties of Ca^2+^-signaling proteins. Our findings thus identify the flexible N-terminal domain as a key determinant of Sorcin activation and a likely general tuning element among penta-EF-hand calcium sensors.

## Methods

### Sequence alignment and conservation analysis

Sorcin protein sequences from nine representative vertebrate species spanning amphibians, reptiles, birds, and mammals were retrieved from the UniProt and summarized in Table S1. Multiple sequence alignments were performed using ClustalW^41^, and pairwise sequence identities and similarities were calculated relative to human Sorcin. Human Sorcin isoforms and splice variants, together with all human penta-EF-hand calcium-binding proteins (sorcin/SRI, grancalcin/GCA, ALG-2/PDCD6, peflin/PEF1, and the calpain small subunit/CAPNS1 and CAPNS2), were aligned by the same procedure to compare N-terminal domain length and composition. Alignments were visualized and annotated using Jalview^42^ with the glycine-rich NTD indicated.

### Protein expression and purification

DNA encoding full-length human Sorcin (SorcinFL; residues 1-198) or the N-terminal deletion construct SCBD (residues 35-198) was cloned into the pET151/D-TOPO expression vector with an N-terminal hexahistidine affinity tag followed by a tobacco etch virus (TEV) protease cleavage site. Protein expression and purification followed protocols previously established in our laboratory^11,27^. Briefly, proteins were expressed in Escherichia coli Rosetta(DE3) cells. Cultures were induced with 0.3 mM IPTG at 16 °C overnight and harvested by centrifugation. Cell pellets were resuspended in lysis buffer (40 mM Tris-HCl pH 7.5, 480 mM KCl, 1 mM EGTA, 2 mM β-mercaptoethanol (βME)) and disrupted by sonication. After removal of insoluble material by centrifugation, the clarified lysate was applied to a nickel-affinity column. Bound protein was eluted with an imidazole gradient, and the affinity tag was removed by TEV protease digestion. Sorcin-containing fractions were further purified by ion-exchange chromatography (mono Q) and size-exclusion chromatography (S75) in 40 mM HEPES-KOH pH 7.5, 150 mM KCl, and 0.5 mM TCEP. Protein purity was assessed by Coomassie staining of SDS-polyacrylamide gel electrophoresis (SDS-PAGE), and concentrations were determined spectrophotometrically at 280 nm using extinction coefficients of 25,900 M^-1^ cm^-1^ for full-length Sorcin and 19,940 M^-1^ cm^-1^ for SCBD.

### Preparation of isotopically labeled proteins

Triple ^2^H/^15^N/^13^C-labeled SCBD and doubly ^2^H/^15^N-labeled full-length Sorcin were expressed in E. coli BL21 cells cultivated in 1 L of M9 medium containing ^13^C-enriched glucose, ^15^N-enriched ammonium chloride, and D₂O (Sigma-Aldrich) as the carbon, nitrogen, and deuterium sources, respectively, as we reported previously^27^. The labeled proteins were purified using the same procedure as the unlabeled proteins, and purity was assessed by SDS-PAGE. All NMR samples were prepared in 90% (v/v) H₂O / 10% (v/v) D₂O at final concentrations of 254 µM for SCBD and 540 µM for full-length Sorcin, in 40 mM HEPES-KOH pH 7.5, 150 mM KCl, 0.25 mM TCEP, and 1 mM EGTA.

### Stopped-flow light scattering

Rapid Ca^2+^-induced aggregation kinetics of full-length Sorcin and SCBD were measured on an Applied Photophysics SX20 stopped-flow spectrometer using Pro-Data SX software, following our previously established protocol^11^. Protein and CaCl_2_ solutions were prepared in matched buffer (40 mM HEPES-KOH pH 7.5, 150 mM KCl, 1 mM βME) and loaded into separate syringes; for each shot, 60 µL from each syringe was driven into the 120 µL observation cell and rapidly mixed. A 565 nm LED served as the light source, and scattered light was detected at 90° by a photomultiplier tube (PMT), with the recorded voltage (V) reporting the extent of aggregation. Measurements were performed at 22 °C, and full-length and SCBD kinetics were compared under identical protein concentration (7 µM) and Ca^2+^ concentration (500 mM) in the same buffer.

### Plate reader turbidity assay

Because SCBD aggregation proceeds on the minute-to-hour timescale, the Ca^2+^-, protein-, and temperature-dependence of SCBD aggregation were measured by a light-scattering (turbidity) assay in a temperature-controlled microplate reader (SpectraMax iD5) in 384-well plates, monitoring scattering at 350 nm. Turbidity is measured as apparent absorbance but reported as scattering. Reactions contained SCBD, CaCl_2_, 40 mM HEPES-KOH pH 7.5, 150 mM KCl, and 1 mM βME; aggregation was initiated by adding CaCl_2_ as the final component. Calcium-concentration, protein-concentration, and temperature-dependence experiments were performed using the ranges specified in the corresponding figure legends. All measurements were performed in triplicate.

### Kinetic analysis

As reported previously^11^, aggregation time courses from the turbidity assay were fit to a double-exponential model to extract kinetic parameters: AU(*t*) = AU_i_ + (AU_m_ - AU_i_)/(1 + 0.5 × exp^kfast × (t1/2 - *t*)^ + 0.5 × exp^kslow × (t1/2 - *t*)^). Here AUᵢ and AUₘ are the initial and maximum scattering signals, t_1/2_ is the aggregation half-time, and k_fast_ and k_slow_ are rate constants. The lag time (τ) was the intercept of the tangent at t_1/2_ with the Vᵢ baseline τ = t_1/2_ - (AU_m_-AU_i_)/(2 × k_1/2_). The maximum rate (kₘₐₓ) and its time (η) were obtained from derivatives of smoothed curves. The dependence of the maximal scattering signal on Ca^2+^ concentration was fit to the Hill equation AU_m_ =AU_max_ × C^n^/(K_1/2_^n^ + C^n^). Here C is the Ca^2+^ concentration, n the Hill coefficient, and K_1/2_ the half-maximal aggregation concentration. For Arrhenius analysis lnk_abs_=-E_a_/R + ln(1/T) and lnk_abs_ was plotted against 1/T, where k_obs_ is the observed rate constant, Eₐ is apparent activation energy, R is the ideal gas constant, and T is the temperature. Because turbidity reports macroscopic particle formation, Eₐ is an apparent activation energy for the overall aggregation process.

### NMR spectroscopy and resonance assignment

NMR spectra were recorded at 37 °C on a Bruker AVANCE III 600-MHz spectrometer with a triple-resonance cryogenic probe (full-length Sorcin) or a Bruker 900-MHz spectrometer (SCBD). Shifts were referenced to trimethylsilylpropanoic acid at 0.0 ppm. SCBD backbone assignments were obtained from HNCACB, CBCA(CO)NH, HNCO, and HN(CA)CO experiments. Data were processed with TopSpin 4.5.0 and NMRFAM-SPARKY and analyzed with CARA. Full-length assignments were taken from BMRB 52475. Comparison ^1^H-^15^N TROSY-HSQC spectra used the same buffer and temperature; field strength and protein concentration differed as specified above. SCBD peaks were matched by residue identity, deposited assignments, and local spectral context.

### Chemical-shift perturbation analysis

Differences between the full-length Sorcin and SCBD spectra were quantified using the combined amide chemical-shift perturbation: Δ*δ* = )(Δ*δ_H_*)^2^ + (0.2 Δ*δ_N_*)^2^ . Here Δδ(^1^H) and Δδ(^15^N) are the shift differences between full-length Sorcin and SCBD; the 0.2 factor scales the broader ^15^N chemical-shift range. CSPs were calculated only for confidently assigned, resolved peaks in both spectra. Descriptive thresholds were set at the mean, mean + 1 SD, and mean + 2 SD after excluding residues 35-42, whose shifts are dominated by the newly created terminus. Peaks present in full-length Sorcin but absent from SCBD, as well as prolines and unassigned residues were denoted in the plot.

### Molecular dynamics simulations

Atomic models of dimeric SCBD were built from the Ca^2+^-bound crystal structure (PDB 4USL)^14^ after deleting all residues preceding Leu35. Missing hydrogens were added with the psfgen plugin in VMD^43^ using the CHARMM36m topology^44,45^, and histidine protonation states were assigned at pH 7.5 based on PROPKA^46^. The model was solvated in a cubic TIP3P water^47^ box (150 × 150 × 150 Å) and neutralized with KCl to 150 mM, giving 325,655 atoms. Systems were minimized and equilibrated with NAMD3^48,49^ and the CHARMM36m force field^44,45^: 5,000 steps of conjugate-gradient minimization followed by 10 ns of MD (1 ns with all protein heavy atoms and Ca^2+^ restrained, 1 ns with only the backbone restrained, and 8 ns unrestrained NPT at 1 atm). Temperature was held at 310 K with a Langevin thermostat (damping 5.0 ps^-1^) and pressure with a Nosé-Hoover Langevin piston barostat^50,51^. Long-range electrostatics used particle-mesh Ewald under periodic boundary conditions^52^, with a 12 Å van der Waals cutoff and a 10-12 Å switching function.

Production runs on Anton 3^34^ were initiated from the final equilibration frame and performed in the NVT ensemble at 310 K using the Antithetic thermostat and Multigrator framework^53^, with a 2.5 fs time step (modified r-RESPA integrator^54^; short-range forces every step, long-range electrostatics every three steps via the u-series/SinhGS method^55^), a 9 Å van der Waals cutoff, and hydrogen bond lengths constrained with M-SHAKE^56^. Frames were saved every 1.2 ns (non-water atoms) and 12 ns (all atoms), and each production simulation was run for 10 µs. To enable a direct comparison with the full-length protein, four 10-µs SCBD simulations were performed using the same protocols as the previously reported full-length Ca^2+^-bound simulations^33^: one unrestrained run initialized in the crystallographic Ca^2+^-bound active conformation, and three runs with Ca^2+^-coordination distance restraints: minimal (single bidentate Glu) at k = 0.05, extensive (all coordinating charged residues) at k = 0.05, and extensive at k = 1 kcal·mol^-1^·Å⁻². To reanalyze NTD-SCBD interactions, we used the previously reported apo full-length trajectory, unrestrained Ca^2+^-bound full-length trajectory, and the three restrained Ca^2+^-bound full-length trajectories from Ye et al^33^.

### Analysis of molecular dynamics trajectories

Global conformational deviation was quantified by C_α_-RMSD relative Ca^2+^-bound reference structure (PDB 4USL)^14^ after C_α_ alignment using residues 35-198. Ca^2+^ coordination and dissociation were monitored as the distances between each Ca^2+^ ion and the carboxylate carbons of its coordinating Asp/Glu residues. An NTD-SCBD contact was present when the minimum residue-pair heavy-atom distance was ≤4.5 Å. Occupancy was the fraction of frames satisfying this criterion. Intra- and inter-protomer contacts were calculated separately and combined for comparison with the protomer-averaged NMR data. Contacts across the covalent NTD-SCBD boundary were excluded. Results were checked with 4.0- and 5.0-Å cutoffs supporting the same conclusion. All data analysis were performed using VMD^43^.

## CRediT authorship contribution statement

### DATA AVAILABILITY

Data will be made available on request.

## Acknowledgments

We acknowledge the technical support provided by the Institute for Bioscience and Biotechnology Research (IBBR) Information Technology Department, particularly Gale Tempest and Scott Carlson. Y.L. and the Liu Laboratory are supported by start-up funding from the University of Maryland, College Park, and the State of Maryland. This project was also supported by an IBBR Seed Grant funded through the University of Maryland Strategic Partnership: MPowering the State. Anton 3 computer time was provided by the Pittsburgh Supercomputing Center (PSC) through Grant 1R24GM154042 from the National Institutes of Health. The Anton 3 machine at PSC is made available by D. E. Shaw Research. The authors also acknowledge the University of Maryland High Performance Computing (HPC) resources (https://hpcc.umd.edu), including the Zaratan HPC Cluster and the IBBR HPC Cluster, which were used to conduct the research reported in this paper.

## Appendix A. Supplementary material

