## Supplemental for "A Conserved Flexible N-Terminal Domain Tunes the Calcium Sensitivity of Sorcin by Stabilizing Its Active Conformation"

Table S1. Summary of protein sequences used for multiple sequence alignment.

| Protein | Gene | Organism | UniProt ID | Length (aa) | Identity* (%) | Similarity* (%) |
|---|---|---|---|---|---|---|
| **Sorcin** | SRI | Homo sapiens | P30626 | 198 | 100.0 | 100.0 |
| **Sorcin** | SRI | Mus musculus | Q6P069 | 198 | 96.0 | 98.5 |
| **Sorcin** | SRI | Canis lupus familiaris | A0A8I3RY81 | 198 | 99.0 | 99.5 |
| **Sorcin** | SRI | Callorhinus ursinus | A0A3Q7P3R5 | 198 | 99.0 | 99.5 |
| **Sorcin** | SRI | Tursiops truncatus | A0A2U3V686 | 198 | 98.5 | 99.5 |
| **Sorcin** | SRI | Pteropus alecto | L5JPW3 | 198 | 98.0 | 99.5 |
| **Sorcin** | SRI | Gallus gallus | Q5ZM28 | 198 | 83.8 | 92.4 |
| **Sorcin** | SRI | Pantherophis guttatus | A0A6P9AIA9 | 195 | 85.4 | 89.9 |
| **Sorcin** | SRI | Xenopus laevis | Q569R9 | 196 | 79.4 | 89.4 |
| **Sorcin** | SRI | Homo sapiens | P30626-2 | 183 | 91.4 | 91.4 |
| **Sorcin** | SRI | Homo sapiens | P30626-3 | 180 | 87.0 | 88.0 |
| **Sorcin** | SRI | Homo sapiens | A0ACI8PYI9 | 170 | 85.9 | 85.9 |
| **Sorcin** | SRI | Homo sapiens | C9J0K6 | 155 | 77.8 | 77.8 |
| **Sorcin** | SRI | Homo sapiens | A0ACI8Q3A7 | 208 | 95.2 | 95.2 |
| **ALG-2** | PDCD6 | Homo sapiens | O75340 | 191 | 36.0 | 53.2 |
| **Grancalcin** | GCA | Homo sapiens | P28676 | 217 | 56.2 | 70.5 |
| **Peflin** | PEF1 | Homo sapiens | Q9UBV8 | 284 | 28.8 | 39.9 |
| **Calpain small subunit 1** | CAPNS1 | Homo sapiens | P04632 | 268 | 26.5 | 38.9 |
| **Calpain small subunit 2** | CAPNS2 | Homo sapiens | Q96L46 | 248 | 26.4 | 43.6 |

*Both sequence identity and similarity were calculated by comparing to the canonical human Sorcin sequence (UniProt P30626).

Table S2. Summary of molecular dynamics simulations.

| Simulation Name | Simulation System | Initial Conformation | Initial $Ca^{2+}$ Occupancy | No. of Atoms | $Ca^{2+}$ Restrains | Spring Constant ($kcal \cdot mol^{-1} \cdot Å^{-2}$) | Simulation Length |
|---|---|---|---|---|---|---|---|
| SIM1 | SCBD | Active | Yes | 325,655 | No | - | 10 μs |
| SIM2 | SCBD | Active | Yes | 325,655 | minimal | 0.05 | 10 μs |
| SIM3 | SCBD | Active | Yes | 325,655 | extensive | 0.05 | 10 μs |
| SIM4 | SCBD | Active | Yes | 325,655 | extensive | 1 | 10 μs |
| SIM0* | Sorcin | Inactive | No | 320,504 | - | - | 10 μs |
| SIM1* | Sorcin | Active | Yes | 324,637 | No | - | 10 μs |
| SIM2* | Sorcin | Active | Yes | 324,637 | minimal | 0.05 | 10 μs |
| SIM3* | Sorcin | Active | Yes | 324,637 | extensive | 0.05 | 10 μs |
| SIM4* | Sorcin | Active | Yes | 324,637 | extensive | 1 | 10 μs |

*Simulations of the isolated Sorcin calcium-binding domain (SCBD) were performed in the present study. Full-length Sorcin simulations marked with an asterisk were reported previously by Ye et al. and were reanalyzed here for comparison.

**A**

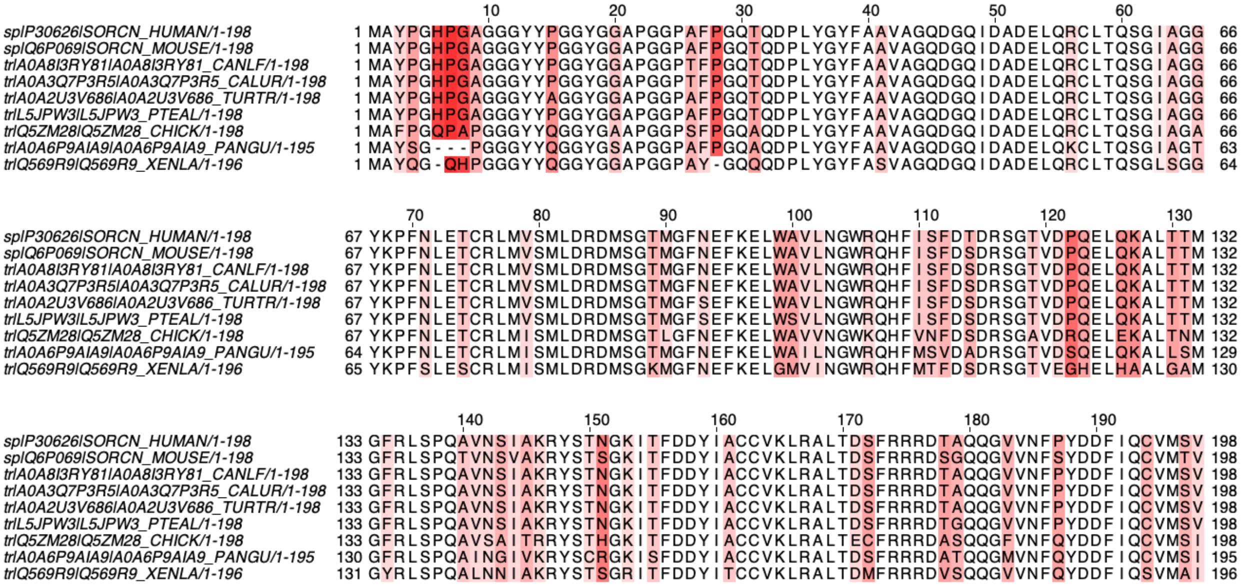

**B**

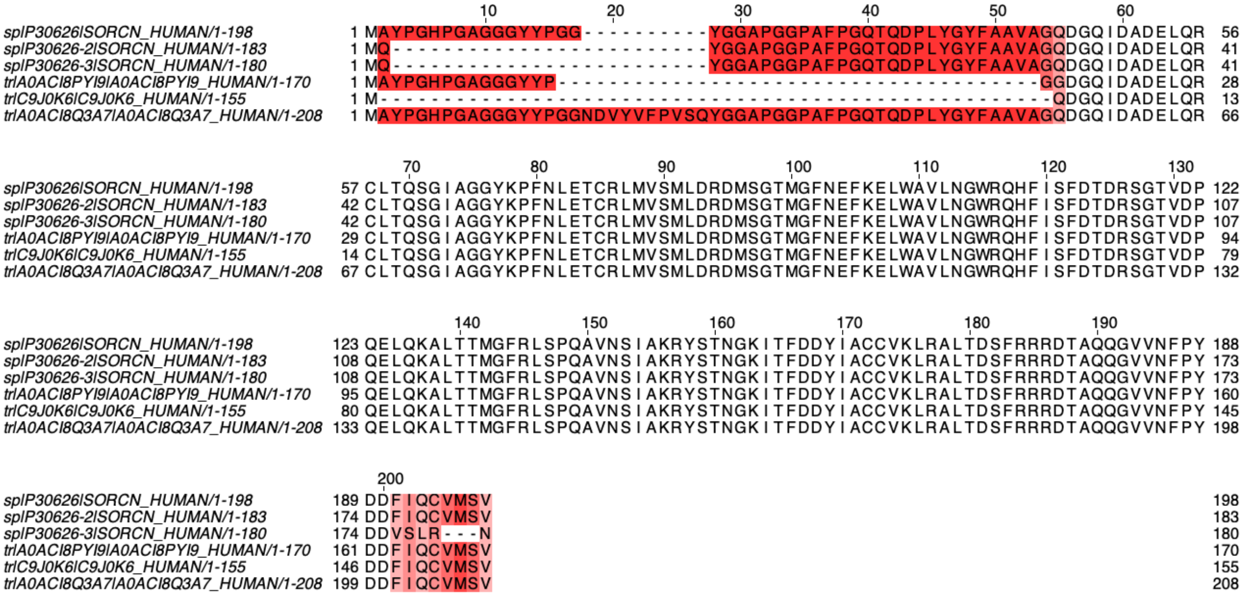

Figure S1. Multiple sequence alignment of Sorcin orthologs and isoforms. (A) Alignment of Sorcin sequences from nine representative species spanning amphibians, reptiles, birds, and mammals. (B) Alignment of six Sorcin isoforms, with the canonical human Sorcin sequence shown at the top.

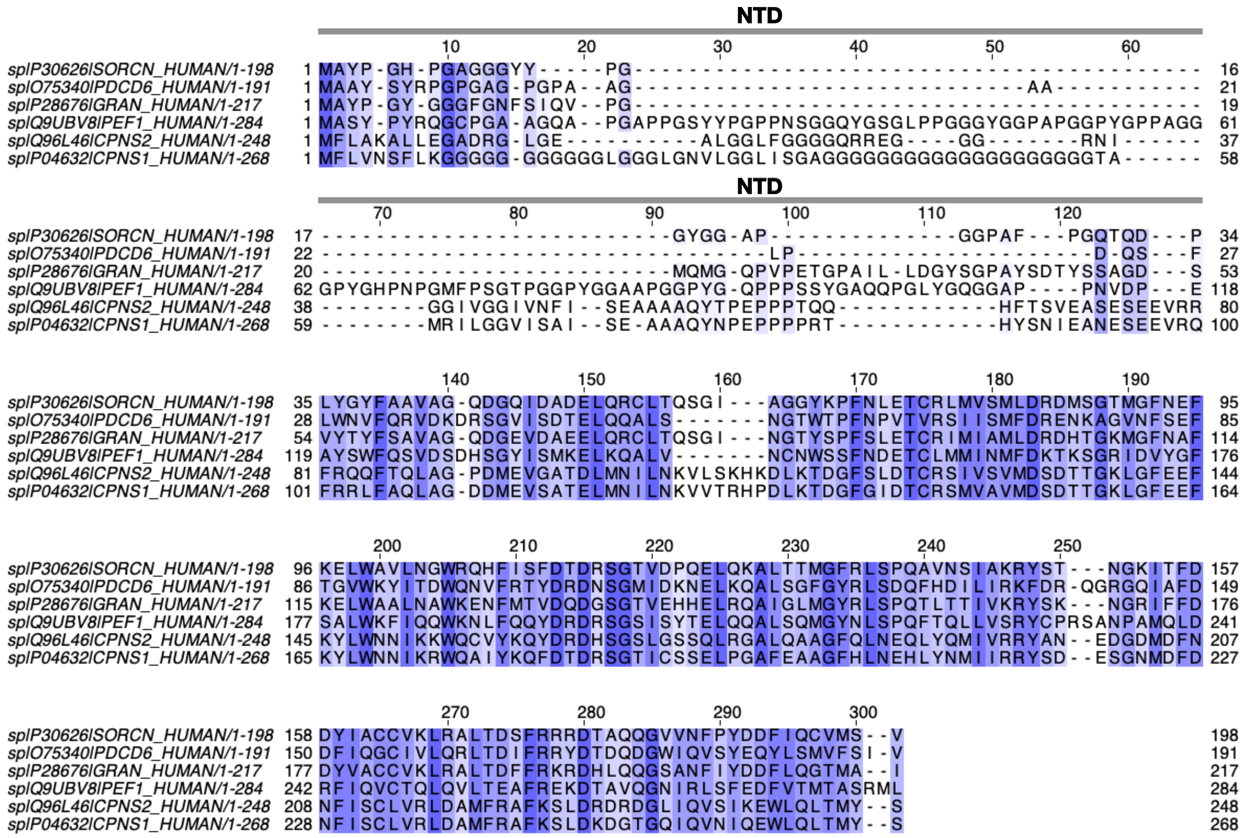


Figure S2. Multiple sequence alignment of the human penta-EF-hand protein family. Conserved residues are highlighted in blue. All family members possess a glycine-rich N-terminal domain (NTD), but the NTD length varies considerably across the family.

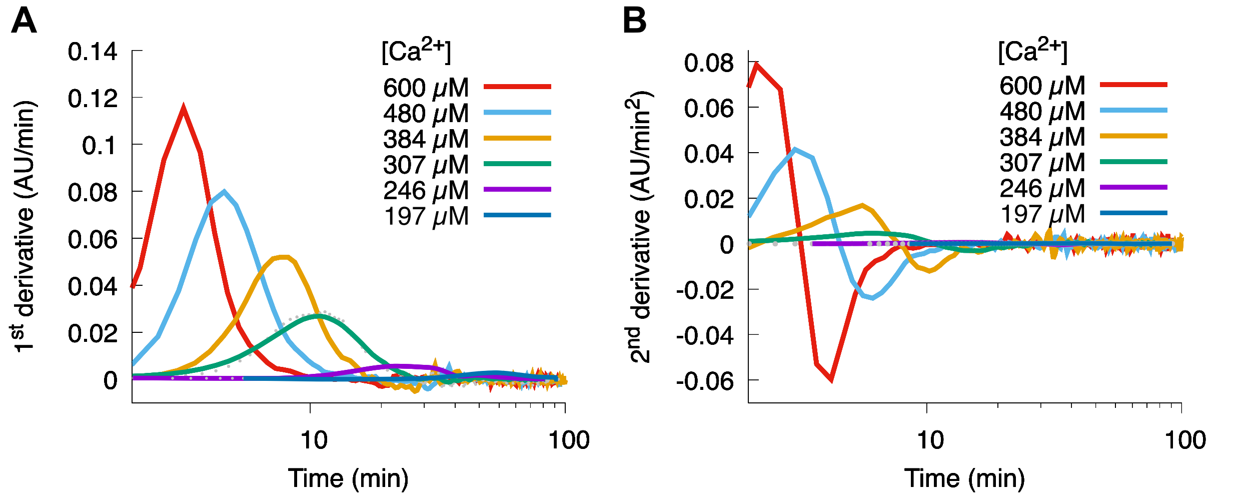


Figure S3. First and second derivatives of representative SCBD aggregation time courses at various $Ca^{2+}$ concentrations. The first (A) and second (B) derivatives of the turbidity curves from Figure 2A, used to determine the maximum aggregation rate ($k_{max}$) and the time at which it occurs (η).

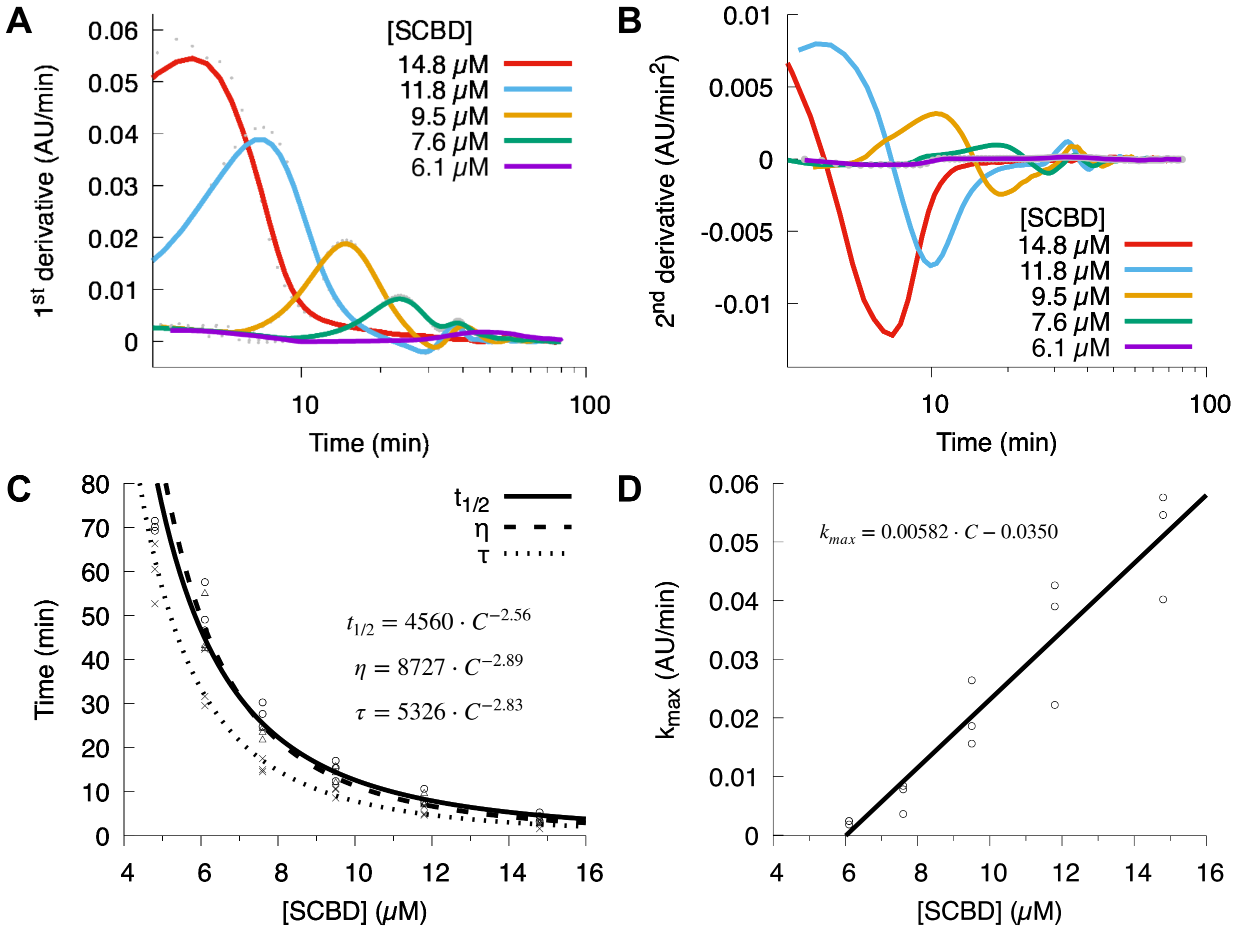


Figure S4. Kinetic analysis of the protein-concentration dependence of SCBD aggregation. First (A) and second (B) derivatives of the turbidity curves from Figure 3A, used to determine the maximum aggregation rate ($k_max$) and the time at which it occurs (η). (C) The aggregation half-time (t½), the time to the maximum aggregation rate (η), and the lag time (τ) as a function of SCBD concentration. Individual values of t½, η, and τ are shown as circles, triangles, and crosses, respectively. (D) The maximum aggregation rate ($k_max$) scales linearly with SCBD concentration (solid line); individual experiments are shown as circles.

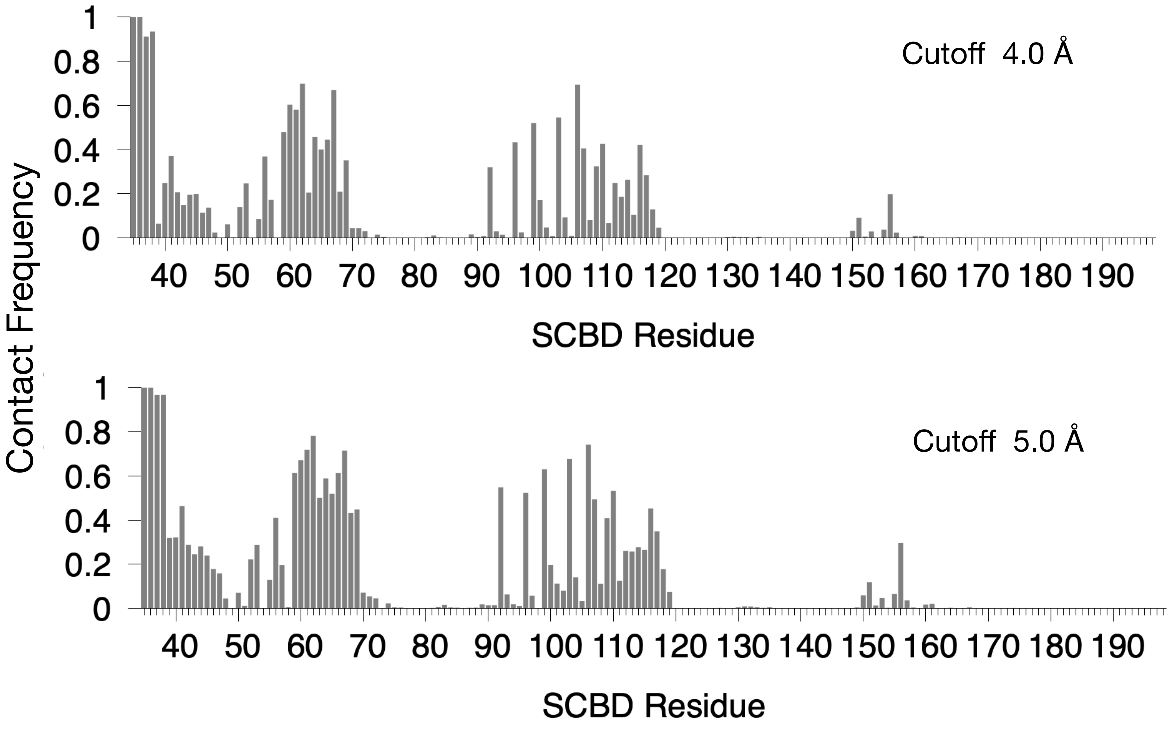


Figure S5. Per-residue contact frequency of the SCBD with the NTD is insensitive to the choice of distance cutoff. For the same trajectory analyzed in Figure 6, distance cutoffs of 4.0 Å and 5.0 Å were tested; both reproduce the contact-frequency pattern shown in Figure 6 (which uses 4.5 Å), differing only in overall magnitude.

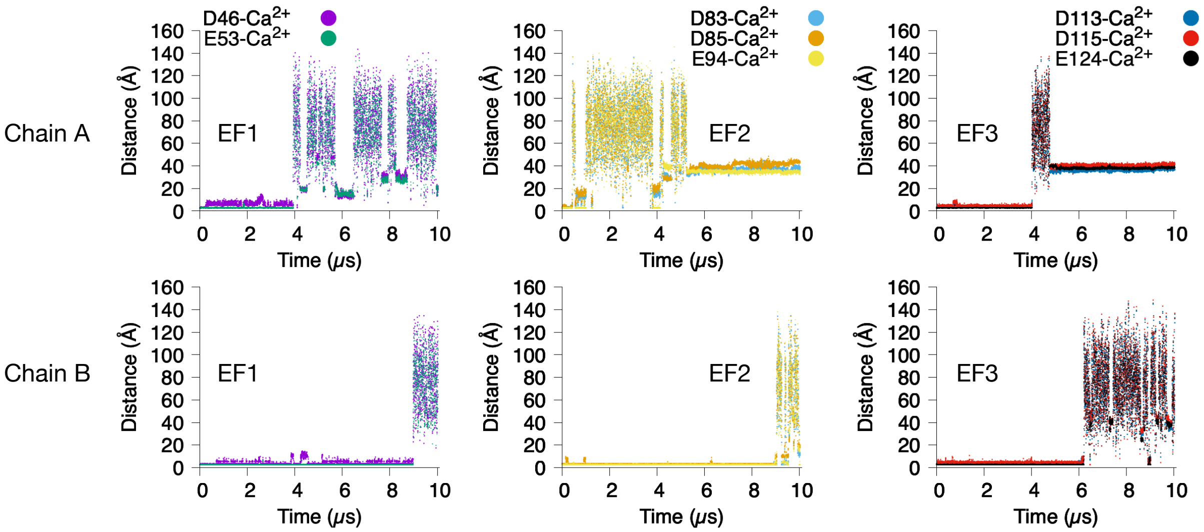


Figure S6. Rapid $Ca^{2+}$ dissociation from the EF-hands in the unrestrained SCBD simulation. Time course of the distances between each $Ca^{2+}$ ion and its charged coordinating residues in EF1–EF3, for the SCBD simulation initiated from the $Ca^{2+}$-bound active state with no distance restraints applied.
